# Group size constraints may mask underlying similarities in social structure: a comparison of female elephant societies

**DOI:** 10.1101/099614

**Authors:** Nandini Shetty, P. Keerthipriya, T.N.C. Vidya

**Author notes:** Corresponding author T.N.C. Vidya Evolutionary and Organismal Biology Unit Jawaharlal Nehru Centre for Advanced Scientific Research Jakkur, Bengaluru – 560 064, India.

## Abstract

We report on female Asian elephant social structure in Nagarahole and Bandipur National Parks (Kabini population), southern India, and examine the role of group size in affecting the outcome of social structure analysis in female elephants, which show high fission-fusion dynamics. Based on five years of data, we found the Kabini association network structured into highly modular communities that we call clans. We then modified the dataset (to obtain the Kabini 500-m dataset) to match sampling methods previously used in a study each of Asian (Uda Walawe) and African savannah (Samburu) elephants, so that network and association statistics could be compared across populations. Measures of association and network structure previously used were more similar amongst the Asian elephant populations compared to Samburu. The Samburu population formed a hierarchically-nested multilevel society whereas the Asian populations did not. However, we found hierarchical clustering levels in all three populations using Louvain community detection. Moreover, the average community sizes obtained through the Louvain method were not significantly different across populations, indicating basic similarities in social structure. Since fission-fusion dynamics allow for community members to form groups of different sizes, we examined the effect of average group size on association and network statistics. Higher average association index and degree, and lower average path length in Samburu compared to the Kabini 500-m dataset were explained by the larger average group size in Samburu. Thus, underlying similarities in the social networks of species showing fission-fusion dynamics may be obscured by differences in average group size.

**Significance Statement:** Various measures of associations and social network analyses have been used to compare social structures of different populations. We studied the social structure of female Asian elephants in a southern Indian population and compared it with those of a Sri Lankan Asian elephant population and an African savannah elephant population. We showed that, while there were social differences between the Asian and African savannah elephant populations using previous methods, there were basic similarities across all three populations using a method of network community detection. This discrepancy across analyses partly stemmed from differences in average group size between populations. Average group size in fission-fusion societies variously affected different association and network statistics, which has implications for inferences about social structure.

## Introduction

Social structure and organization, which include the patterning of relationships and the system of interactions between individuals, may affect foraging, reproductive opportunities, anti-predatory benefits, vulnerability to disease, and information transfer (e.g. 1-8), making them important in the study of animal species. Social organization is thought to evolve in response to resource-risk distributions (9-14), and one of the modal types of mammalian social organization was called the fission-fusion society, in which groups fuse together or split away in response to spatio-temporally varying resources, balancing the costs and benefits of group-living (e.g. 13, 15-20). Distinct types of fission-fusion societies were identified: multilevel societies that were either strictly hierarchically nested (e.g. hamadryas) or flexibly nested (e.g. gelada baboons), and the classical or individual-based fission-fusion society (e.g. chimpanzees and spider monkeys) (see 21). It has since been recognized that fission-fusion societies form a continuum of different extents of fission-fusion dynamics (see 22). Here, by analyzing social structure in female elephants, we show how group size may bridge modal fission-fusion societies. Group size is the number of individuals sighted together in the field and may often be smaller than the size of the mostinclusive, socially-meaningful community in species showing fission-fusion dynamics.

Female elephants show high fission-fusion dynamics (see 22), and previous studies have suggested a multitiered (hierarchically-nested multileveled) social structure in African savannah elephants (19, 23) and non-nested, multileveled social structure in an Asian elephant population (24). The differences between these social structures may arise from group size limitation in the Asian elephant, preventing hierarchical structure from being apparent, but this has not been examined previously, as only one detailed study of Asian elephant social structure (25) was available. Since observed social structure may reflect evolved patterns, as well as plastic responses to the current environment (see 26, 27), studies of multiple populations are required to understand the social structure of a species. Here, we examine the role of group size in affecting social structure by collecting the first large-scale quantitative data on Asian elephant social structure from India, from the Nagarahole-Bandipur (Kabini) population, and by comparing this with data from the Uda Walawe Asian elephant population in Sri Lanka, and the Amboseli and Samburu African savannah elephant populations, for which published data on female social structure are available (19, 23-25, 28-29).

The Asian elephant (*Elephas maximus*) is an endangered species, whose social organization may have been impacted to varying extents across its range by prolonged, historic manipulation by humans. Therefore, there have been calls for detailed studies from multiple populations in order to understand the drivers of their social organization (24-25, 30-31). Asian and African savannah elephants form matriarchal societies, with females and their dependent offspring living together in groups, and adolescent males dispersing from the groups and leading largely solitary lives thereafter (23, 31-34). However, based on previous studies, there seemed to be differences in social structure within and between elephant species, possibly from different sampling methods and ecology (see below).

The African savannah elephant exhibits a multitiered female society (19, 23, 32), with mother-offspring units being the basic units, and “family group” referring to one to a few closely related mother-offspring units (35). In Amboseli, family groups identified at the beginning of the study were later called core groups, and associations of family or core groups were termed bond groups (23, 28). Family groups that shared dry-season home ranges were called clans (23). Social tiers in Samburu were identified statistically through cluster analysis and included, hierarchically, second-tier units (family groups), third-tier units (kinship groups of Douglas-Hamilton (32) or bond groups of Moss and Poole (23)), and fourth-tier units (19). Groups themselves were differently identified in the field, with individuals of core groups having to be within 100 m of one another in order to qualify as being associated in Amboseli (28), and individuals within a 500-m radius of an aggregation centre being classified as a group in Samburu (19). Amboseli and Samburu have also experienced different extents of poaching (36-37), but the association network in Samburu was found to be resilient to the elevated levels of poaching (38). Samburu and Amboseli are similar in elephant density and ecology (39) and social tiers are similar in the two populations (Table 1). Therefore, differences between the female social networks of the two populations, with the Samburu network being much more interconnected than the Amboseli network (Fig. 1), are likely to stem from differences in sampling methods.

**Table 1.** Details of social tiers in previously studied elephant populations and from the present study in Kabini. Clan sizes in Kabini are based on 16 clans seen at least 40 times each. The other data come from Fernando and Lande (30) for Ruhuna National Park (Sri Lanka), de Silva *et al.* (25) for Uda Walawe (Sri Lanka; the values were calculated using Louvain community detection from data kindly provided by Shermin de Silva), Moss and Poole (23), Lee (78), and Moss and Lee (47) for Amboseli (Kenya), and Wittemyer *et al.* (19) and Goldenberg *et al.* (38) for Samburu (Kenya) (except for the cell marked ‡, whose values were calculated using Louvain community detection from data in 24, kindly provided by the authors).

| Population | Ave. no. of females in a family group (range) | Ave. no. of individuals in a family group (range) | Ave. no. of females / family groups in a bond group | Ave. no. of individuals in a bond group (range) | No. of females in a clan / fourth-tier unit / most inclusive unit | No. of individuals in a clan / fourth-tier unit / most inclusive unit |
| --- | --- | --- | --- | --- | --- | --- |
| Kabini | – | – | – | – | Mean: 13.31, SD: 7.78, Median: 11.5, IQR: 7.5-18.5, Max: 32 | Mean: 29.19, SD: 19.76, Median: 21, IQR: 17-39.5, Max: 83 |
| Kabini 500m, ≥20 sightings | – | – | – | – | Mean: 15.57, SD: 9.74, Median: 18, IQR: 9-19.5, Max: 31 | – |
| Uda Walawe | – | – | – | – | Mean: 11.67, SD: 7.47, Median: 12, IQR: 5-17, Max: 23 | – |
| Ruhuna | – | – | – | – | 7.75 (4-11)<sup>†</sup> | 14.75 (7-24)<sup>†</sup> |
| Amboseli | 2.35 (1-9)* | 7.22 (2-23)* | 2-5 family groups | – | 5-9 family groups | Range: 50-250 |
| Samburu | 2.2* (1-5) | 7.64 (1-15) | 2.0 family groups (4.4 females on ave., based on *) | 16 (6-40) | ‡Mean: 13.75, SD: 7.46, Median: 11, IQR: 8.8-16, Max: 28 | Median: 33.5, IQR: 28.8-80.3; Median: 32, IQR: 23.5-38 during a subsequent period. |

\*Family groups in Amboseli comprised, on average, 2.35 females and 7.22 total individuals in 1976, while core groups (erstwhile family groups that had expanded) in 2002 comprised an average of 7.08 females (47). The former are shown so that appropriate comparisons can be made across populations. †These were described as family groups originally (because families contained up to 10-20 females and their offspring according to Wilson (60)), but Asian elephant populations are placed in the last two columns here as the most inclusive social units. IQR: Inter-Quartile Range.

**Fig. 1.**
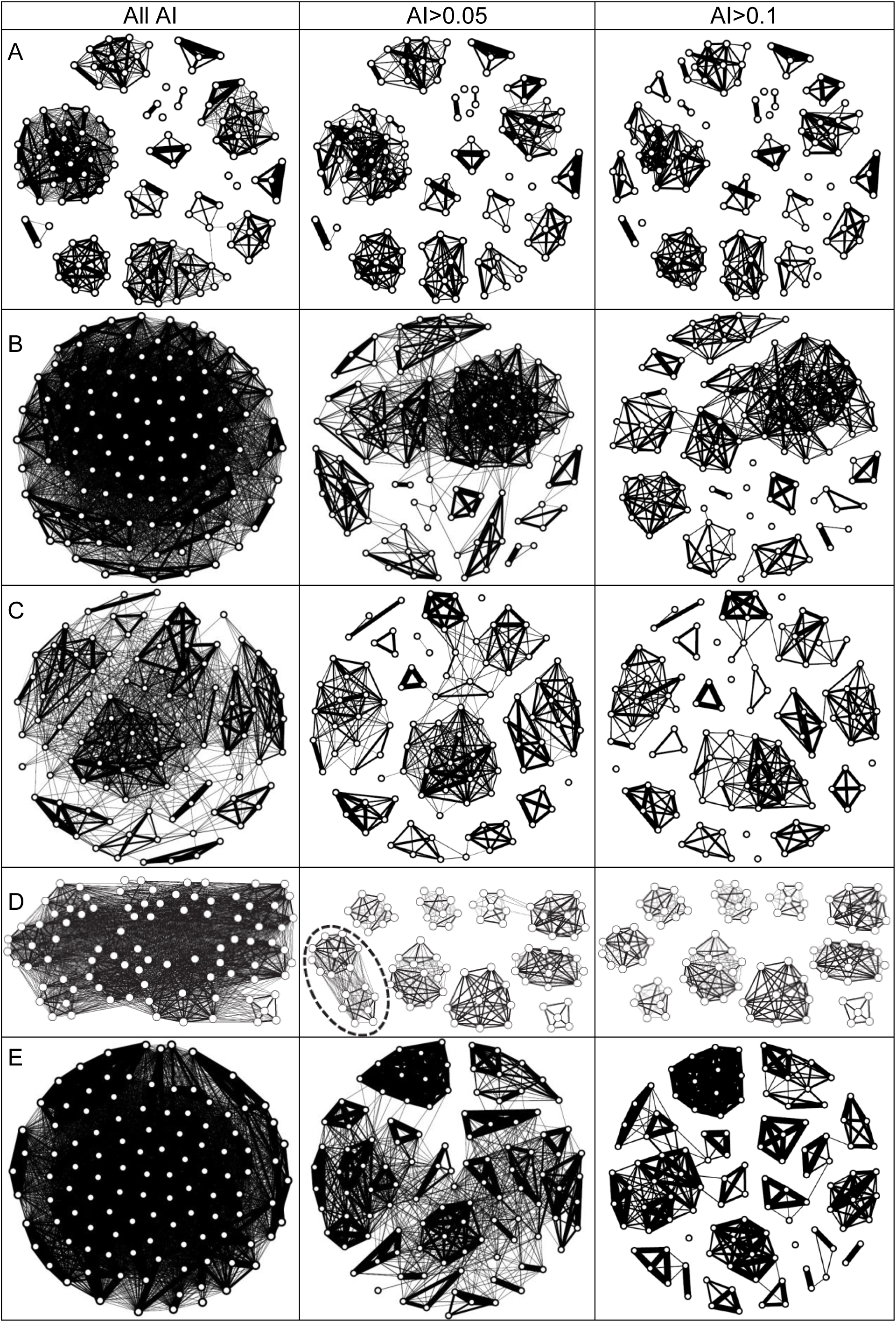
Social networks in A) Kabini, original dataset, B) Kabini 500-m dataset, C) Uda Walawe (Sri Lanka), D) Amboseli (Kenya), and E) Samburu (Kenya), based on all associations (first column), associations with an AI cutoff of 5% (second column), and an AI cutoff of 10% (third column). The networks in C and E are based on de Silva and Wittemyer (24; data kindly provided by the authors) and those in D, from Archie and Chiyo (29; reproduced with the permission of the publisher, John Wiley and Sons, license number 4025511083875). The dashed oval in D indicates a bond group. Only individuals sighted at least 20 times are included in the Kabini networks, as was the case in the Uda Walawe and Samburu networks.

Studies on female Asian elephant social organization had suggested a matriarchal society with fission-fusion dynamics, inferred from female social groups of varying sizes (33-34). However, the precise nature of social organization was ambiguous, with studies from Sri Lanka largely not describing multitiered societies but those from southern India implying them (see below). McKay (33), in southeastern Sri Lanka, described the most inclusive female social group as a “herd” (of 15-40 individuals), which could contain subunits that showed fusion and fission, but which did not associate with other “herds” that shared their home range. Fernando and Lande (30) found smaller group sizes subsequently (see Table 1), but these groups too did not associate with other groups that shared their home range, and were referred to as family groups. A study in southern India (34) suggested the existence of a multitiered society, with “family groups” (single adult female and her dependent offspring), “joint-family groups” (two or more adult females and their offspring), “bond groups”, and “clans” (50-200 individuals). Baskaran *et al.* (40), in southern India, referred to social associations of females that showed coordinated movement and were presumably related as a “clan” (of up to 65 individuals), but did not demarcate social tiers within clans. The first large, quantitative study of Asian elephant social organization, carried out in Uda Walawe, Sri Lanka, found female social organization with long-term associates, and larger social units than typically seen associating at any time in the field (25), which was also the case with the previous, less quantitative studies (and indeed expected in fission-fusion societies in general). However, unlike the previous studies, the larger social units (“herds” or “family groups” or “clans” as the term might be) were connected to one another in a social network at the level of the entire population (24-25). In a comparative analysis, female groups in Uda Walawe were smaller, showed weaker associations, and were less connected at the population level than those in the Samburu African savannah elephant population (24-25). The Uda Walawe population thus showed a non-nested, multilevel society, with individuals associating differently with two types of social affiliates, in contrast to a multitiered society in Samburu with nested social tiers (24). Although some of the initial confusion relating to female Asian elephant social organization seems to have stemmed from an attempt to equate social levels in the Asian elephant with those described in the better-studied African savannah elephant, there was also the possibility of female Asian elephant social organization being different between Sri Lanka and the mainland because of extensive historical disturbance to elephants in Sri Lanka compared to southern India (pp. 68-69 in 41, 42).

We used data from the Nagarahole-Bandipur (Kabini) population in southern India to find out whether female social structure in this population was similar to that in Sri Lanka, and, if so, whether the difference in social structure from that of the African savannah elephant could be explained by a constraint on group size. We also wanted to find out whether the wider social network in Uda Walawe compared to associations found in previous studies in Sri Lanka could have resulted from differences in methods. A 100-m distance cutoff had been traditionally used to delineate Asian elephant groups (30, <150m in 33), while a 500-m cutoff, similar to the one in Samburu, had been used in Uda Walawe (25). We expected that there might be lower levels of connectedness in the Uda Walawe population compared to the Kabini population because of extensive historical disturbance in the former. However, on the whole, we expected greater similarity between Kabini and Uda Walawe, with smaller group sizes and lower network connectivity in the Asian elephant populations than in the African savannah elephant because of ecological differences. We further attempted to find out how group sizes would affect network statistics in general, given these fission-fusion societies.

## Results

The dataset used to examine female associations comprised 3893 sightings of female groups, sighted between 2009-2014, in which all the females that were ten years old or older (referred to simply as females in the rest of the paper) could be identified. These 3893 sightings constituted 87% of all the female group sightings during the study period, and comprised 9551 individual female sightings, including repeat sightings of the same individuals. A group was an aggregation of female elephants that showed coordinated movement and/or behaviours, and were usually within 50-100 m of one another. Members of a group were said to be associating with one another. The number of uniquely identified females from this dataset was 330. Since we wanted to compare our data with the Uda Walawe and Samburu populations in which a 500-m distance cutoff had been used to identify associations, we created an additional dataset (the Kabini 500-m dataset) in which we grouped together females that were within 500 m of one another.

### Association network and AI in the Kabini population based on the original dataset

Data on female group sightings were used to calculate an Association Index (AI, see Methods) between each pair of females. AIs were used to construct association networks. The association network based on the original Kabini dataset showed clearly demarcated communities (Fig. 2), with associations between females being highly non-random compared to Poisson expectation (*G*-test for goodness of fit, *G*=1514.46, *df*=23, *P*<0.001) and based on permutation tests (Table S3). The overall network modularity, which is a measure of the extent to which a community is partitioned, was high (0.936). Community structure within networks was identified using the Louvain method (43, see Methods). We refer to the final communities obtained as clans, in keeping with previous terminology used to refer to the most inclusive female social grouping of elephants in southern India. The largest clan in our study consisted of 32 females (83 individuals, including their offspring). We did not find female associations across clans during over five years, except on seven occasions (Fig. 1). Upon executing the Louvain algorithm (and excluding the 30 single females that were rarely seen), we found 60, 40, and 39 (which corresponded to the eventual 39 clans) communities after the first, second, and third rounds (passes) of hierarchical community detection, respectively. Thirteen of the 39 clans showed two levels (i.e., changed in composition from the first to the second pass) and one clan showed three levels of social organization (changed from the first to the second, and from the second to the third passes), whereas the remaining clans showed a single social level (did not change across passes). The clans that showed more than one social level were significantly larger (average=14.4, SD=7.57, *N*=14) than those that had a single social level (average, SD, *N* of clan size: 3.9, 2.74, 25; Mann-Whitney *U* test: *U*=20.0, *Z*_adj_=-4.598, *P*<0.001, Fig. S4). Based on clans that were sighted over 150 times, we found that 95% of the clan members were sighted on average within the first 40 sightings of the clan (and 92% within the first 30 sightings). Since under-sampling could lead to incomplete clans, we additionally analyzed only clans sighted >40 times and found that they were also significantly larger when they were composed of more than one social level (average=17.0, SD=7.29, *N*=10) compared to a single level (average=7.2, SD=3.66, *N*=6; Mann-Whitney *U* test: *U*=6.0, *Z*_adj_=-2.609, *P*=0.009, Fig. S4).

**Fig. 2.**
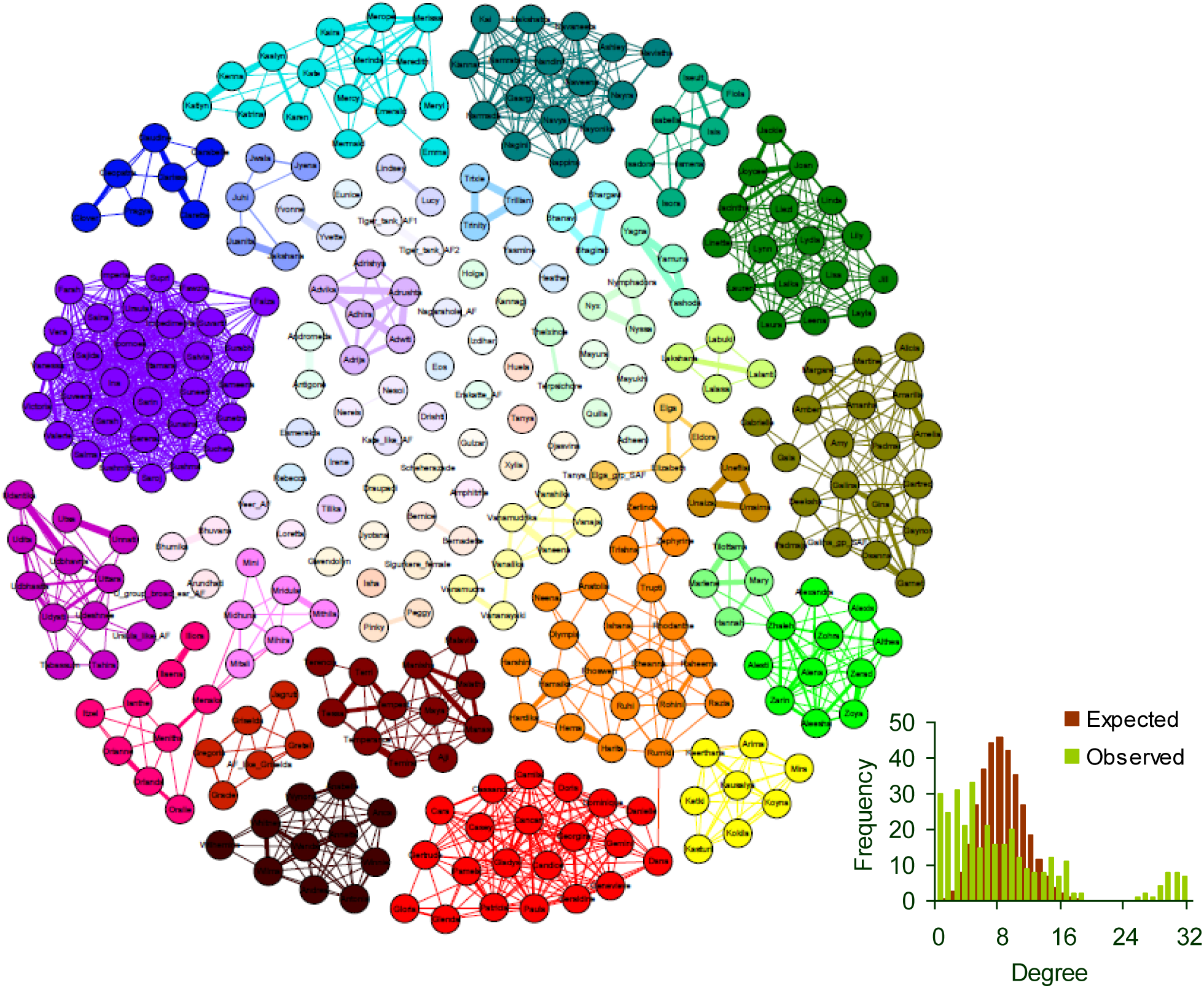
Social network based on the entire dataset of 330 females drawn using the Fruchterman Reingold layout (77) in Gephi. Each node here is a female and the edges between the nodes indicate nonzero AI between females (edge thickness is proportional to AI). Nodes are coloured based on modularity classes and we refer to nodes of the same colour as a clan. The expected (Poisson) and observed degree distributions based on this social network are shown. The average degree was 8.32 based on this network, which includes individuals seen only once; when such individuals were removed, the average degree was 9.55 (274 individuals). Most of the solitary nodes towards the centre are females that were seen only once or a few times. The small number of connections across clans arose from seven sightings during the five year study period. Four of these were due to associations between a female with a newborn calf (that could not keep pace with the group) with other clan females.

In keeping with clearly defined clans with few associations between them, the overall AI distribution was highly skewed (Fig. 3), with only 2.5% of the AI values being non-zero (average AI=0.004, SD=0.040, median=0) and 10.8% being non-zero when only individuals that were seen at least 20 times were included (Table 2). The average degree (number of associates of a female) and average distance-weighted reach (a measure of a female’s reach in the association network, see Methods) were low (Table 2) because of female associations being restricted to clans. The average clustering coefficient (the probability that two randomly chosen neighbours of a focal female are connected, see Methods) was high due to the large number of connections within clans, and density, which measures connectedness across the entire network, was low (Table 2).

**Fig. 3.**
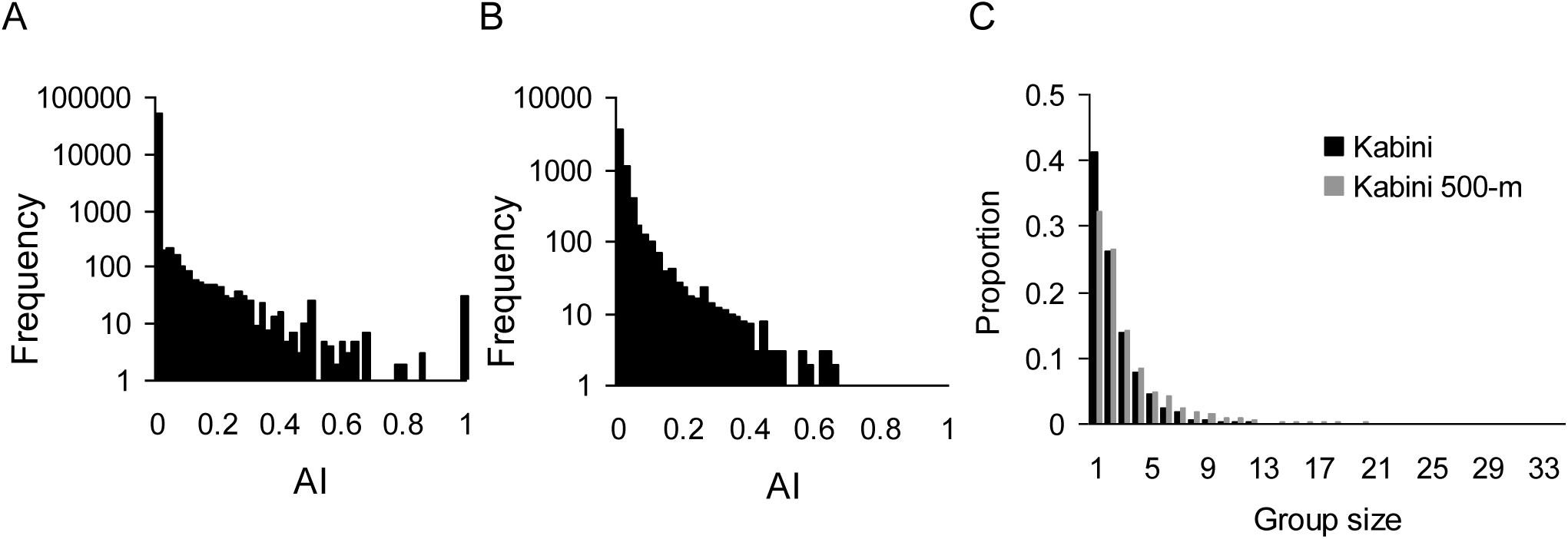
AI distributions based on A) the entire original Kabini dataset and B) the Kabini 500-m dataset, and C) group size distributions based on the original Kabini dataset and Kabini 500-m dataset. The relatively high frequency of AI=1 in A) is because of small number of sightings of some individuals, and this disappears if only females seen at least 20 times are considered.

**Table 2.**
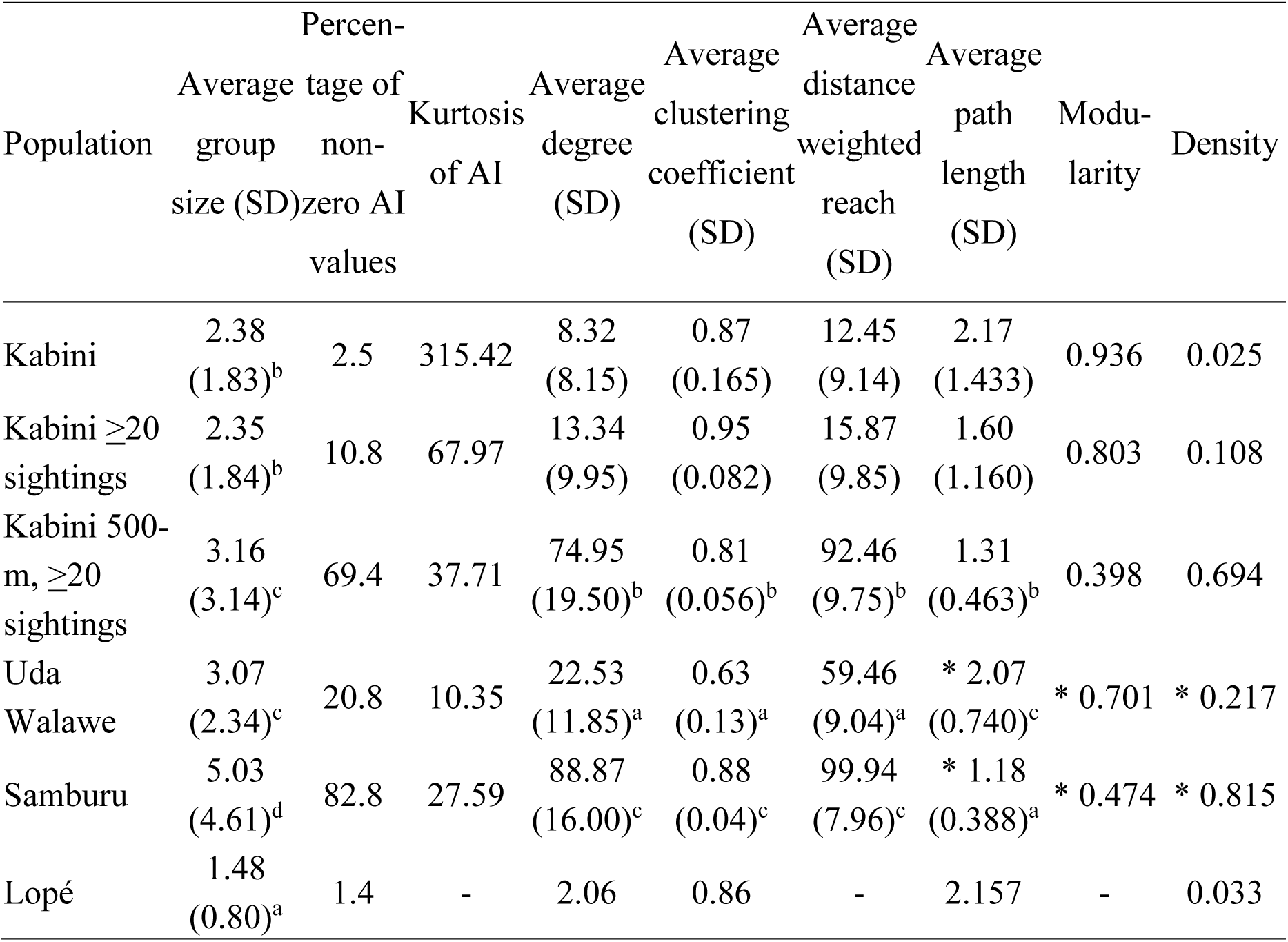
Average group size (number of females), AI statistics, and network statistics for different elephant populations. Statistics for the Uda Walawe and Samburu populations are reproduced from de Silva and Wittemyer (24) and the ones with asterisks were calculated from network files (of 24) kindly provided by Shermin de Silva and George Wittemyer. Statistics for the Lopé forest elephant population are reproduced from Schuttler *et al.* (44). Statistics such as the degree and density might be underestimates in Lopé because the number of times individuals were sighted was very small (network statistics based on individuals sighted at least twice) and there was a significant correlation between the number of sightings and the number of associates (44). The small average group size is, however, in keeping with that found in forest elephants in Dzanga Bai (average female group size including dependants: 2.7, SD=1.3) in a long-term study (51). The average group size for Kabini ≥20 sightings is the average of group sizes of only those sightings in which all the females were seen at least 20 times (this is shown for the sake of completeness). Significant differences in metrics based on the Welch’s two-sample test are shown using superscripted alphabets (a<b<c), with *α* set to 0.0017 based on a flat Bonferroni correction for 29 tests.

### Comparison of association networks across populations

The association network based on the original Kabini data was highly disconnected, unlike female social networks in the previously studied African savannah elephant and Uda Walawe Asian elephant populations (24, Fig. 1, first column), but more connected than the network in the Lopé African forest elephant population (44). However, since different criteria had been used for grouping females, we compared the Kabini 500-m network with the Uda Walawe and Samburu networks, and the original Kabini network with the Amboseli network (in which associations had been recorded somewhat similarly, see 28). The Kabini 500-m networks were intermediate in connectedness between the Samburu and Uda Walawe networks (Fig. 1), with the average degree, average distance-weighted reach, and average clustering coefficient in the Kabini 500-m network being significantly smaller than those in Samburu (Welch’s two-sample tests: average degree: *U*=5.772, *df*=208.3, *P*<0.001, average distance-weighted reach: *U*=6.216, *df*=207.9, *P*<0.001, average clustering coefficient: *U*=10.636, *df*=195.3, *P*<0.001), but significantly larger than those in Uda Walawe (average degree: *U*=23.862, *df*=179.3, *P*<0.001, average distance-weighted reach: *U*=25.687, *df*=211.7, *P*<0.001, average clustering coefficient: *U*=13.068, *df*=140.2, *P*<0.001; Table 2). The average path length (the number of connections on the shortest path between two females) in the Kabini 500-m network was also intermediate, being larger than that in Samburu (Welch’s two-sample test: *U*=16.573, *df*=11452.6, *P*<0.001) and smaller than that in Uda Walawe (*U*=64.999, *df*=9038.0, *P*<0.001). Visually, the network based on our original data was less connected than that of the Amboseli population when there was no AI cutoff, but similar to Amboseli at AI cutoffs of 5% and 10% (Fig. 1).

The original Kabini network did not change substantially when an AI cutoff of 0.05 was used, unlike networks from all the other datasets (Fig. 1). The Kabini 500-m network changed dramatically at an AI cutoff of 0.05 like the Samburu network. However, network structure curves, which illustrate the cohesiveness of social networks at different association strengths (see 25, Methods), of the Kabini 500-m and the Uda Walawe datasets were roughly similar in shape and differed from that of the Samburu population (Fig. 4, see 24). The AI distribution based on the Kabini 500-m dataset (Fig. 3) bore a greater visual similarity to that of Uda Walawe than to that of Samburu, as high AI values were absent (see 24) and this similarity in AI distribution could have given rise to the similarity in network structure curves. The average AI was, however, significantly smaller in Uda Walawe (0.019) compared to that in Kabini 500-m dataset (0.034; Welch’s two-sample test: *U*=11.195, *df*=11295.3, *P*<0.001), which was in turn significantly smaller than that in Samburu (0.049; Welch’s two-sample test: *U*=8.209, *df*=9544.9, *P*<0.001). However, the percentage of non-zero AI values was much higher in the Kabini 500-m dataset than in Uda Walawe (Table 2). The kurtosis of the Kabini 500-m dataset was higher than those of both Samburu and Uda Walawe (Table 2).

**Fig. 4.**
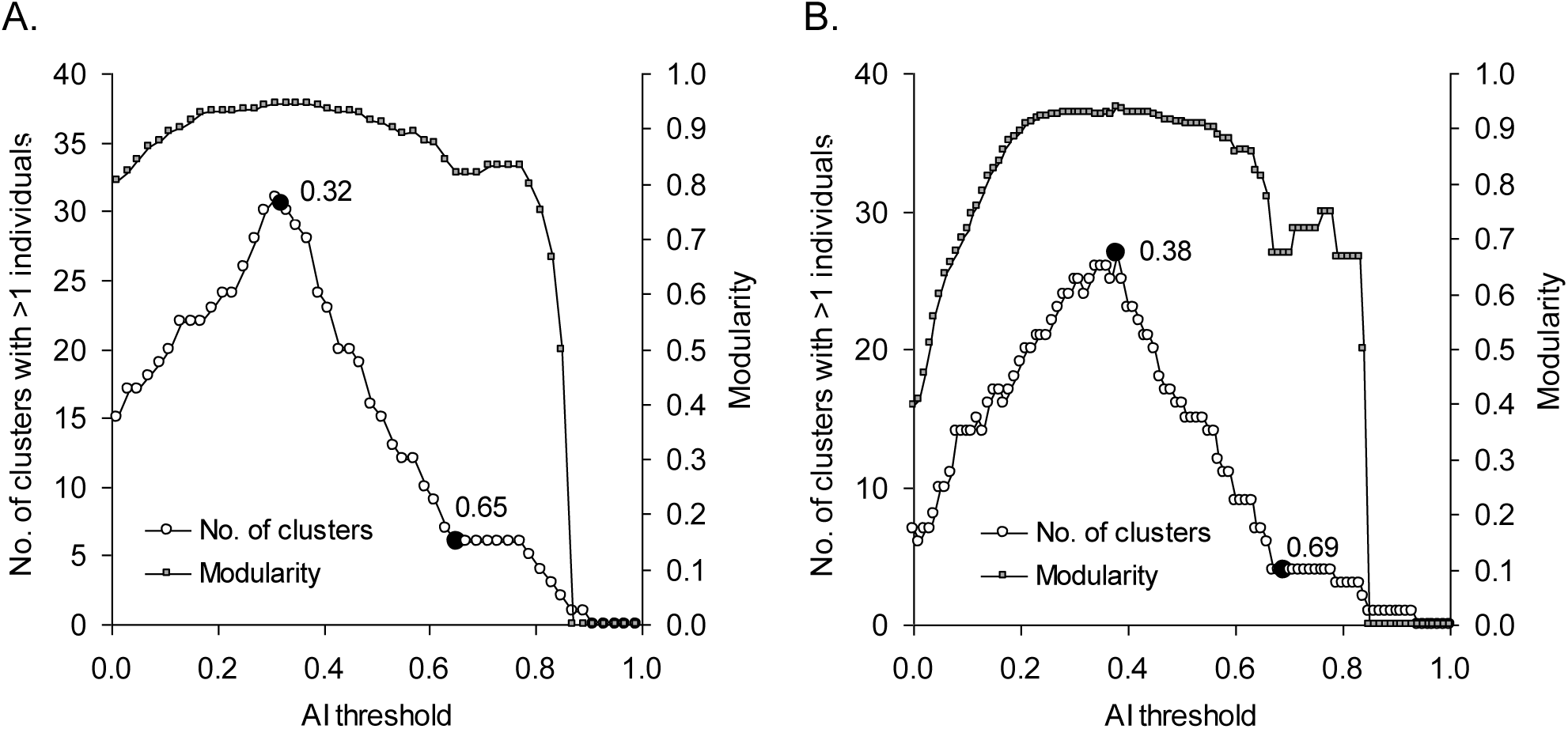
Network structure curve (of females seen at least 20 times) for A) the original data and B) data based on a 500-m distance cutoff, showing two points of slope change based on window of 0.3 (*P*<0.001 for AI threshold of 0.32 and *P*=0.006 for AI threshold of 0.65, points of slope change were 0.33 and 0.665 based on a window of 0.2; *P*=0.001 for AI threshold of 0.38 and *P*=0.012 for AI threshold of 0.69, *P*=0.082 for the AI threshold of 0.69 based on the 0.2 window).

Louvain community detection on the Kabini 500-m dataset, and Uda Walawe and Samburu datasets (from 24, kindly provided by the authors) showed up to two social levels in the Kabini 500-m and Uda Walawe datasets, and up to three levels in the Samburu dataset (although sometimes, Uda Walawe showed up to three levels and Samburu, up to two, see SI 5). The numbers of communities after the first pass were 20, 16, and 24 in the Kabini 500-m, Uda Walawe, and Samburu, datasets, respectively, and the numbers of communities after the second pass were 7, 9, and 9, respectively. Eight communities were detected after the third pass in Samburu. In the Kabini 500-m dataset, five of the seven final communities changed from the first to the second pass, while the other two remained compositionally the same. The numbers of communities that changed in composition from the first to the second pass were four out of nine in Uda Walawe, and seven out of nine in Samburu. As in the original Kabini dataset, communities with two social levels were larger than those with a single level, although this was not statistically significant in the Uda Walawe dataset (Mann-Whitney *U* tests: Kabini 500-m: *U*=25.5, *Z*_adj_=-2.480, *P*=0.013; Uda Walawe: *U*=39.5, *Z*_adj_=-1.856, *P*=0.063; Samburu: *U*=15.0, *Z*_adj_=-3.808, *P*<0.001). Interestingly, community sizes at a particular community detection round were not different across populations (average±SD after the first pass: Kabini 500-m: 5.45±3.05, Uda Walawe: 6.56±4.87, Samburu: 4.58±2.34, Kruskal-Wallis test: *H*_2,69_=0.380, *P*=0.827; average±SD after the second pass: Kabini 500-m: 15.57±9.74, Uda Walawe: 11.67±7.47, Samburu: 12.22±7.60, Kruskal-Wallis test: *H*_2,25_=0.594, *P*=0.743; Fig. 5A; Mann-Whitney U tests, Kruskal-Wallis test, and test for homogeneity of slopes below carried out using Statistica 8, (45)). There was a correlation between community sizes after the second pass and the number of first-level communities within those second pass communities (Fig. 5B), and a test for homogeneity of slopes showed no difference in slopes across the three populations (Multiple *R*^2^=0.765, *P*<0.001, Effect of population: *F*[2,19]=0.502, *P*=0.613; Effect of the number of first level communities: *F*[1,19]=52.608, *P*<0.001).

**Fig. 5.**
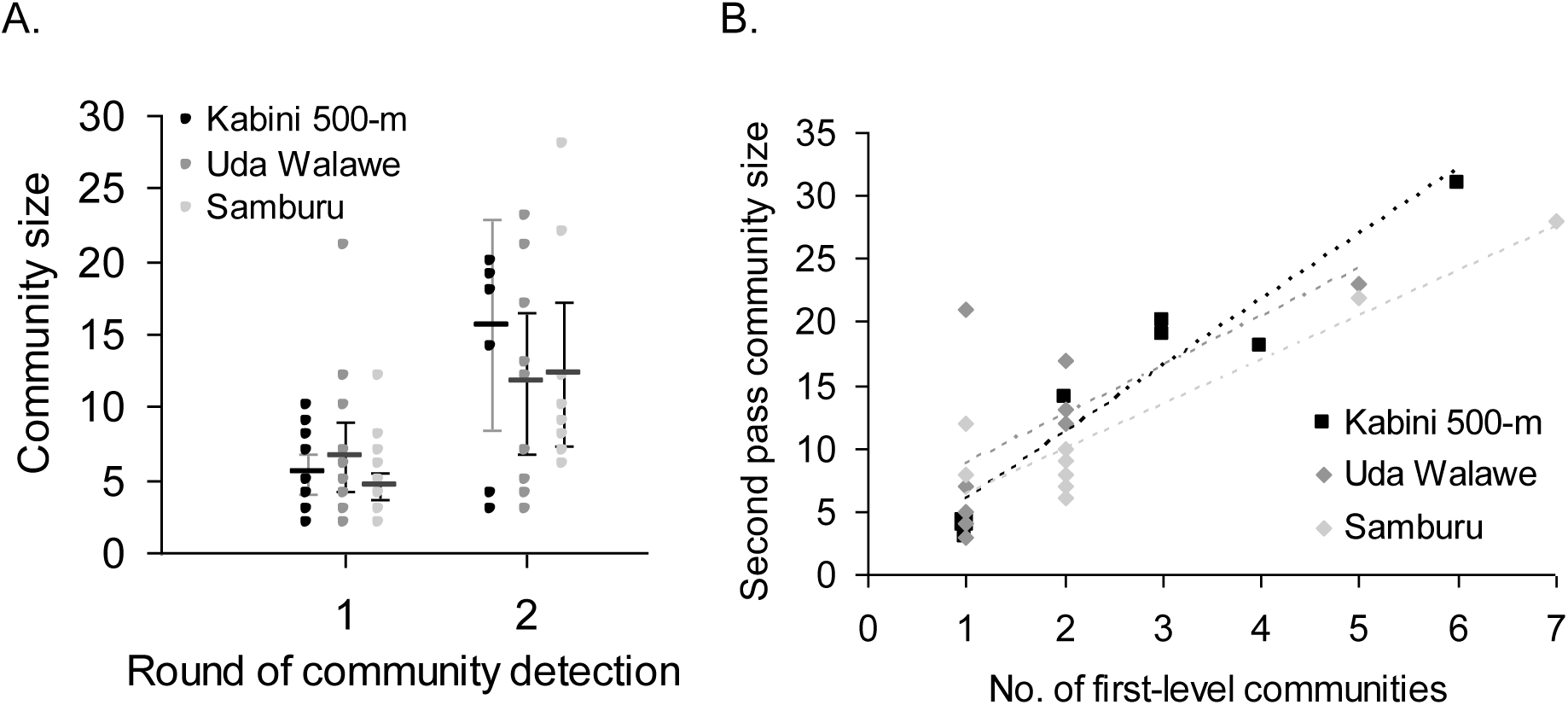
A) Community sizes after the first and second rounds of community detection using the Louvain algorithm, based on the Kabini 500-m, Uda Walawe, and Samburu datasets. Means and 95% CI (1.96 SE) are also shown. B) Sizes of communities after the second pass of community detection, composed of varying numbers of first-level communities in the Kabini 500-m, Uda Walawe, and Samburu datasets. Equations of the trendlines (in the same colours as the data points of the respective populations) for the three populations are: Kabini 500-m: *y*=5.227*x*+0.636, *R*^2^=0.904, Uda Walawe: *y*=3.861*x*+4.803, *R*^2^=0.453, Samburu: *y*=3.521*x*+2.833, *R*^2^=0.859.

### Cluster analysis and cumulative bifurcation curve

We carried out average-linkage clustering of females based on AI and plotted the cumulative number of bifurcations at different linkage distances in order to compare the shapes of the curves across populations. The cumulative bifurcation curves were concave-up in the Kabini and Kabini 500-m datasets (Fig. S6), indicating a smaller number of linkages at small linkage distances (tight associations) than at large linkage distances (loose associations). This was similar to the pattern seen in Uda Walawe and unlike that seen in Samburu, in which the curve was convex (24).

### Observed group sizes and the effect of group size on AI and network statistics in random networks

The average group size in the Kabini population was small, with 2.38 females per group (Table 2, median=2) and the group size distribution was skewed to the right (Fig. 3C). We found Lopé to have a significantly smaller average group size than that in Kabini, the Kabini 500-m dataset and Uda Walawe to have similar group sizes, and Samburu to have a significantly larger group size than the Kabini 500-m and Uda Walawe datasets (Welch’s two-sample tests, Table 2). Since differences in group sizes are likely to affect AI and network statistics, we examined the effect of group size on these statistics, to find out whether differences across populations in these statistics could simply be a result of differences in group size. This was done using random datasets with group size distributions mimicking real distributions by adjusting beta distribution parameters (*α*=1, *β*=7, maximum group size=19 for Uda Walawe, *α*=2, *β*=9, maximum group size=26 for Samburu, and *α*=1, *β*=9.5, maximum group size=27 for Kabini 500-m dataset) (see Methods). Maximum group size was altered to change the average group size. With the exception of some values at very small group sizes, the three different beta distributions of group sizes did not significantly change the expected value of the AI or network statistic considered under random association (Fig. 6). Under all three beta distributions of group sizes, the average expected AI increased linearly with increasing average group size, the average degree and average clustering coefficient increased and plateaued with increasing average group size, and average path length decreased with increasing average group size (Fig. 6).

**Fig. 6.**
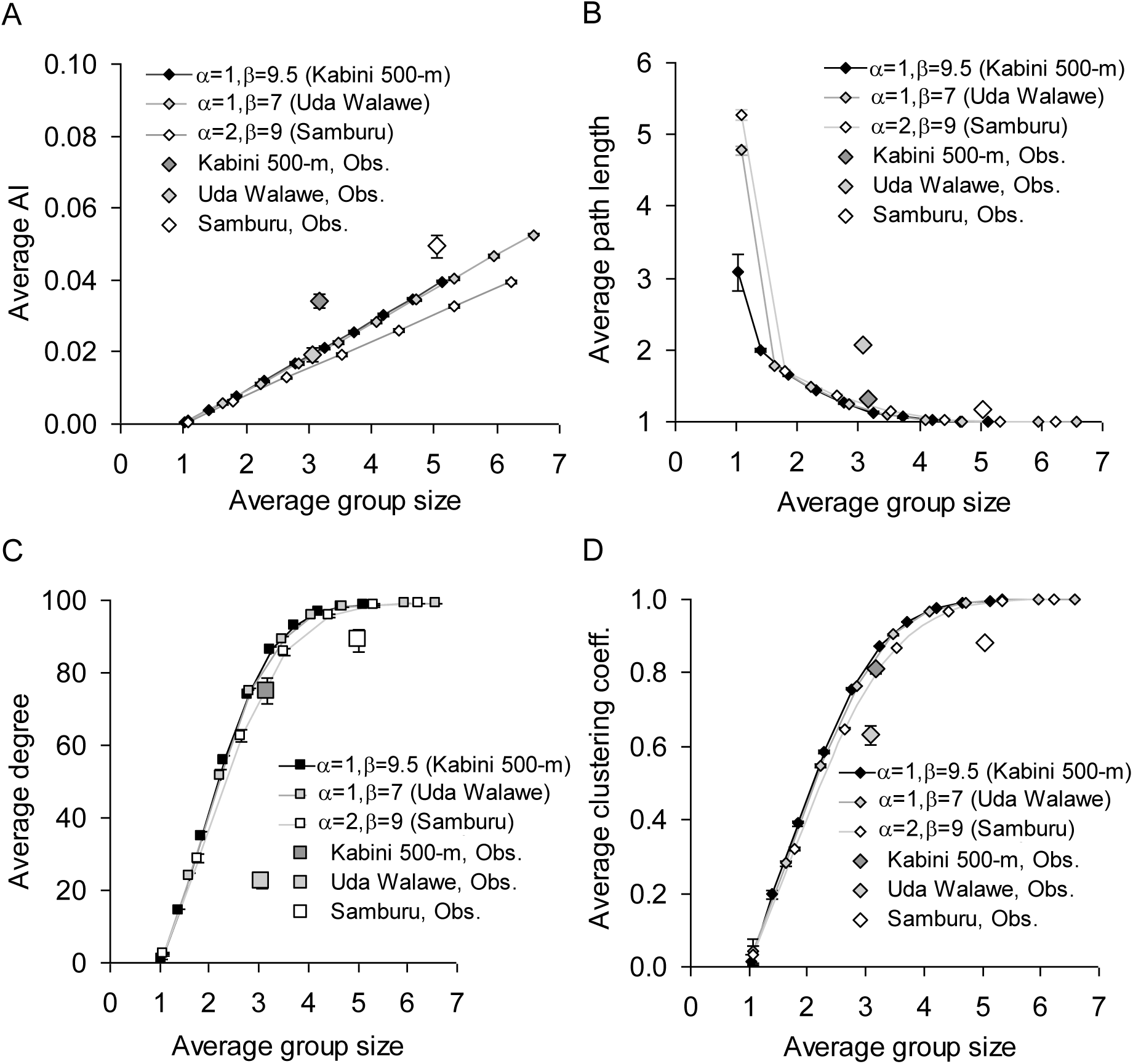
Observed average AI and network statistics from the Kabini 500-m, Uda Walawe, and Samburu datasets, and expected values of these statistics at different average group sizes, based on random datasets with different beta distributions (*α*=1, *β*=9.5; *α*=1, *β*=7; *α*=2, *β*=9, see Methods) of group sizes. A. average AI, B. average path length, C. average degree, and D. average clustering coefficient. All the error bars are 1.96 SE.

For each observed statistic (calculated from field populations with an observed average group size), we calculated an expected random value of the statistic, and obtained an interval of [expected - lower 95% CI of observed]/expected and [expected - higher 95% CI of observed]/expected values for each population. If these intervals overlapped across populations, it indicated that the populations differed from the random expectations to the same extent and, therefore, differences in the statistic could be explained by differences in observed average group size (see Methods). The higher average AI in Samburu compared to the Kabini 500-m dataset could be explained as an effect of group size, with the average AIs being similar when average group sizes were taken into account. The observed average AI in both Samburu and Kabini 500-m datasets were larger than the expected average AI for the corresponding average group sizes to the same extent ([*E-O*]/*E*=-0.667, interval: -0.578 to -0.756 for Kabini 500-m, [*E-O*]/*E*=-0.626, interval: -0.522 to -0.730 for Samburu). The smaller average degree in the Kabini 500-m dataset compared to Samburu could also be explained as an effect of group size differences in the two populations, as the observed average degrees in both these datasets were smaller than the expected average degrees for the corresponding average group sizes to the same extent ([*E-O*]/*E*=0.111, interval: 0.154 0. 067 for Kabini 500-m, [*E-O*]/*E*=0.092, interval: 0.122-0.061 for Samburu). The higher average path length in the Kabini 500-m network than in Samburu (Table 2) could also be explained by differences in average group size (Table S7). We had previously found the clustering coefficient in the Kabini 500-m network to be significantly smaller than that in Samburu (see above), but the former was 5.5% smaller than expected for a random network of the same average group size while the latter was 11% smaller than expected for its average group size. Therefore, corrected for group size, the Kabini 500-m dataset would have a significantly higher clustering coefficient than the Samburu population, although this difference was small (Table S7). Whereas differences between Samburu and Kabini 500-m datasets could largely be explained by differences in average group size, the higher average AI and higher average degree in the Kabini 500-m data compared to those in Uda Walawe (average AI: [*E-O*]/*E*=-0.014, interval: 0.085 to -0.113; average degree: [*E-O*]/*E*=0.722, interval: 0.750-0.694) remained as average group sizes were not significantly different between these two populations.

## Discussion

### >Social structure in the Kabini population

Based on the first quantitative data on social structure of female Asian elephants from India, we found highly non-random associations between females, with the association network being clearly demarcated into communities that we call clans. That there were only seven associations between clans over five years suggests that the clan is the most inclusive level of meaningful social structure in the Kabini population. Using the Louvain method of community detection, we found up to three hierarchical social levels within clans. However, there was variability in clan structure, with 38% of the clans seen more than 40 times showing a single social level, 56% showing two levels, and a single clan showing three levels. The hierarchical levels did not show up in the form of a typical nested multitiered organization (with clear ‘joint-family groups’, bond groups, and clans seen in the field, which seem to have been inferred based on animals sharing a common area, see 34) because of possible constraints on group size. It is not clear whether single social levels in some clans arose from recent permanent fission, demographic factors (see *Implications for Asian elephant social structure* section below), or from clans not being fully identified. While the last is possible, it is not likely as we used a 40-sighting cutoff for clans (as we had found, based on clans sighted over 150 times, that 95% of the clan members were sighted on average within the first 40 sightings of the clan and 92% within the first 30 sightings). It would be interesting to examine the attributes, other than clan size, of clans showing different hierarchical levels in order to understand the differences in clan structure.

### Comparison of social structure across populations and the role of group size

Asian and African savannah elephants were initially thought to share largely similar social organizations (34, 46). Subsequently, they were found to differ in their social structure (24), with larger groups and stronger associations within and across groups in the African savannah elephant. The Kabini 500-m dataset showed intermediate average degree, average distance-weighted reach, clustering coefficient, and path length, compared to those of Samburu and Uda Walawe. The network structure curve, cumulative bifurcation curve, and AI distribution from the Kabini 500-m dataset were more similar to those from Uda Walawe rather than Samburu (see 24). However, contrary to previous finding that kurtosis was higher in African savannah than in Asian elephants (24), we found that the kurtosis of the Kabini 500-m dataset was higher than those of both Samburu and Uda Walawe populations (Table 2). Since kurtosis measures the heaviness of the tail compared to the normal distribution, this result reflects the difference between the average AI and AIs in the tail of the distribution, and not the latter alone (which was highest in Samburu). Visual comparison of the original Kabini dataset’s network with the Amboseli network showed a more connected network in the Amboseli population with no AI cutoff, but similar networks in Kabini and Amboseli at AI cutoffs of 5% and 10% (Fig. 1). However, in the absence of access to the Amboseli network data (29), we were not able to make any further comparisons.

Despite the above differences between the Samburu African savannah elephant and Asian elephant populations, we found through Louvain community detection that there was hierarchical structuring within social networks in the Kabini 500-m dataset, Uda Walawe, and Samburu populations. Since the cumulative bifurcation curve combines data from across the clustering dendrogram, variation across social units in AIs and unequal tiers across social units do not allow for hierarchical structure to be detected (also see 24), which the Louvain algorithm can recover. The number of hierarchical communities found were similar across populations. Although Samburu often showed a third round of clustering (Uda Walawe sometimes showed a third round and Kabini did not), this only resulted in a minor change, with nine communities grouping into eight after the third pass. Community sizes were not significantly different across populations, both after the first and second pass of community detection. There was also no difference across populations in the relationship between second pass community sizes and the number of first-level communities within second pass communities (Fig. 5B). Results from these analyses suggest basic similarities in social structure across elephant species. We show that some of the differences in social structure arose from differences in group sizes across populations. We found that the higher average AI and higher average degree in Samburu compared to the Kabini 500-m dataset arose from different average group sizes in the two populations. Average path length was also similar when corrected for group size. The clustering coefficient in Samburu was smaller than expected at the corresponding group size compared to that in the Kabini 500-m dataset, but the extents to which they differed was small. We used 1.96 SE as the 95% confidence intervals for comparisons of the observed and expected values. It is likely that the errors and, therefore, the overlaps in statistics between populations would actually be larger. Therefore, the tests are conservative, and it is plausible that group size differences account for more of the social structure differences than we find.

Although there is a continuum of societies showing fission-fusion dynamics (22), if they had to be compared to the modal multilevel organizations (see 21) based on AI, network statistics, and cumulative bifurcation curves, the social structure of the African savannah elephant would correspond to the flexible nested society (21; with the lower social level stable and the higher level flexible) or lie between the strict nested and flexible nested multilevel societies, while that of the Asian elephant would correspond to the classical fission-fusion society (with the lower level flexible and the higher level stable) or lie between the classical and flexible nested fission-fusion society. Nestedness does not seem to be complete in the African savannah elephant since partial or whole core groups may associate together to form a larger unit, and single females, although rare (47), have a choice of associating or not with their family group members (29, 47). African forest elephant social organization has been previously compared to the individual-based fission-fusion society of chimpanzees (44). However, as mentioned above, we find underlying similarities in network structure between the African savannah and Asian elephant populations and the differences in network statistics seem to emanate from group size differences. Since the average size of first-level Louvain communities is similar to the average group size in Samburu, individuals of a first-level community can potentially be part of the same group, resulting in high AI values, and easily detectable nestedness. This also results in lower-level social units such as the family/bond group being stable units (see 19). On the other hand, the average sizes of first-level communities are about twice the average group sizes in Uda Walawe and Kabini (Kabini 500-m dataset). When group sizes are restricted, only subsets of the first-level community can associate together, resulting in lower AIs, unstable lower-level social units, and a less nested appearance. This would suggest that the multilevel social structure observed in the Asian elephant is a derived condition due to restricted group size (see ‘Route A’ of Aureli *et al.* (22)), compared to that observed in the African savannah elephant. It is interesting that the average first- and second-level community sizes were not different across elephant populations, indicating that there might be something fundamental about these sizes. It is possible that demographic processes (e.g. 48) or cognitive factors (e.g. 49) influence these community sizes.

The difference in group sizes between the Samburu population and the two Asian elephant populations probably relate to differences in ecology, and more specifically to food resource distribution. Asian elephants typically inhabit moister, more forested habitat than African savannah elephants, and possibly face more challenges obtaining food. African forest elephants, which inhabit wetter and denser habitats, with ephemeral and patchily distributed resources (50), than the Asian elephant on average, show even smaller group sizes (44, 51, see Table 2), a highly disconnected association network, and possibly an individual-based fission-fusion society (44). Turkalo and Fay (52) suggested that, apart from the patchy distribution of food, low predation by humans might explain the small group sizes of African forest elephants compared to African savannah elephants that have faced high poaching pressure. However, despite differences in predation between Sri Lanka (historic human depredation, no current animal predator) and southern India (tigers can prey on calves), there was no difference in the average group sizes in Kabini (Kabini 500-m dataset) and Uda Walawe.

That group size and social structure are interlinked has been obvious (see 53). Grouping patterns modulate interaction opportunities, thus resulting in the social structure seen (22, 27, 53-55). However, we show, for the first time, how social structures uncovered by AI and network statistics in fission-fusion societies may differ primarily because of group size differences. Thus group size differences may mask underlying similarities in the social structures of related species showing fission-fusion dynamics, which can be uncovered by hierarchical community detection.

### Implications for Asian elephant social structure

Despite broad similarities, there were also some differences in Asian elephant social structure based on the limited detailed comparison between one Sri Lankan (Uda Walawe) and one southern Indian (Kabini 500-m dataset) populations. The average number of associates, average association strength, and social network connectedness was greater in the Kabini 500-m dataset compared to Uda Walawe, although average group sizes were not different between the two populations (Table 2). It is possible that the lower levels of cohesiveness in the Uda Walawe population arose from extensive historical disturbance in Sri Lanka, with thousands of elephants having been hunted and captured during the 1800s and early 1900s (see 33, 42), and the elephant population being decimated to only about 1500 individuals by the mid-1900s (see 33). By one estimate, at least 17,000 elephants were hunted, exported, or died in captivity during the 19^th^ century, changing the behaviour and demographics of elephants on the island (see 42). Hunting and capture of elephants in southern India appears to have occurred on a much smaller scale, with no decline in population size (41, pp. 68-69). Moreover, the *kheddah* method used for capturing female elephants in southern India (including in part of our study area, Nagarahole National Park) was intended to capture entire groups rather than isolated individuals (41, pp. 70-73). Therefore, female social organization in southern India is unlikely to have been as impacted as that in Sri Lanka.

Although unrelated females from decimated surviving groups are known to associate together in elephant populations subject to anthropogenic mortality (39, 56-58), breakdown of social structure itself may or may not change group size. Smaller family groups were found in highly poached African savannah elephant populations (see 56, 59), and heavy poaching was also thought to have reduced associations and affected social network structure in Lopé, Central Africa (44). Decimation of the population may increase group size where the habitat allows it, but if there is a resource-based constraint on group size, network cohesiveness is likely to decrease (because of associates being killed) while the group size may not change. We think that this might be the likely scenario in Uda Walawe. Recent anthropogenic disturbance appears to be similar in Kabini and Uda Walawe, with dams built in the late 1960s-early 1970s, submerging forest land and creating reservoirs that elephants now use.

As mentioned previously, associations in the Uda Walawe population had been defined using a 500-m cutoff (24-25), which is why we used the Kabini 500-m dataset for an appropriate comparison. If this was not used, as in our original network, the Uda Walawe network would also presumably be even less cohesive and consist of small communities of females. This might explain the observations of Fernando and Lande (30), who had used a 100-m cutoff to identify groups in southeastern Sri Lanka, and suggested that female social organization was limited to the family level, based on family sizes of 10-20 individuals in African savannah elephants mentioned by Wilson (60). We suggest that the most inclusive level of female social structure, which is also the most stable level, in Asian elephants be called clans. The numbers of females in the most inclusive level in the Kabini population were similar to those in Samburu (Table 1). We also found structuring within some clans, although this is not easy to discern because of groups being small. A common property of clans in the Asian elephant seems to be the lack of association with other clans, despite proximity (“herds” of McKay (33), “family groups” of Fernando and Lande (30), this study). The extent to which the small number of observations resulted in smaller community sizes in Ruhuna is not clear (herd sizes found by McKay (33): 15-40 individuals, by Fernando and Lande (30): 7-24 individuals, by us: inter-quartile range: 17-40, maximum: 83 individuals). The smaller the group size, the longer is the study period required to observe sufficient associations between individuals in order to interpret social structure in a species showing fission-fusion dynamics. However, there were some small clans in Kabini too, arising from deaths of females and/or a series of male offspring, who do not contribute to clan size. Although most of the single females in Fig. 2 are from clans whose ranges are probably at the periphery of our study area, such that we have not yet sighted other females from those clans, there were a few clans that were sighted many times but continued to show a small number of females. The most notable of them included only two females and their dependant offspring, despite being sighted over 300 times. Although a clan of two might as well be called a family group, we prefer to retain the term clan for the most inclusive grouping because, in clans that show structuring, the clan and not the family group seems to be the most stable unit.

## Methods

### Field data collection

The field study was carried out in Nagarahole National Park and Tiger Reserve (Nagarahole; 11.85304°-12.26089° N, 76.00075°-76.27996° E, 644 km^2^) and the adjacent Bandipur National Park and Tiger Reserve (Bandipur; 11.59234°-11.94884° N, 76.20850°- 76.86904° E, 872 km^2^), in the Nilgiris-Eastern Ghats landscape in Karnataka, southern India (Fig. S1). The greater landscape holds the single largest population of over 8,500 (61) Asian elephants in the world, of which about 2,600 (62) elephants probably use Nagarahole and Bandipur. The area sampled was centred around the Kabini reservoir and extended into the forests of Bandipur and Nagarahole, and we refer to the population as the Kabini population (see 63).

Field data were collected from March 2009-July 2014, on a total of 878 field days. Sampling during 2010 was limited because of field permit issues. We drove along pre-selected routes from ~6:30 AM to 6:00-6:45 PM (depending on daylight hours and field permits) and identified female elephant “groups” as aggregations of female elephants, usually along with their young, that showed coordinated movement (especially towards or away from a water source or salt lick), coordinated behaviours (such as bunching and facing the same direction when perceiving a threat from other elephants or heterospecifics), or affiliative behaviour, and were usually within 50-100 m of one another. During our original data collection, we did not use a 500-m distance cutoff because it was clear from the uncoordinated, and sometimes aggressive, interactions between different aggregations of elephants within 500 m of one another that they did not belong to a single social group. Sighting details of elephant groups, including group size, time of sighting, and GPS location were recorded. Individuals were photographed and identified based on multiple natural physical characteristics, and aged based on body size, skull size, and body characteristics, using the Forest Department’s semi-captive elephants of known ages in the area as a reference (see 63). Although individuals older than 15 years have previously been referred to as adults (34, 63), since we subsequently found that females were often sexually mature at 10 years of age (as in other elephant populations, see 64-65), we analyzed associations for females that were 10 years old or older (referred to simply as females in this paper).

### Association data

We retained only sightings in which all the females could be identified. We considered sightings of the same group to be independent if they were observed again after 2.5 hours because this interval yielded roughly similar probabilities of groups either changing in composition or not (Fig. S2). Changes in group composition within this time period were not recorded as separate sightings. In the Kabini 500-m dataset, we grouped together females that were within 500 m of one another, based on GPS data. In this dataset, sightings sharing a common female during the day were merged together into a single sighting, after the manner of de Silva *et al.* (25). Further, only females sighted at least 20 times were retained in the dataset (*n*=109 females) as had been done in the Uda Walawe and Samburu datasets. AI between pairs of females was calculated as the ratio of the number of times two females A and B were seen together (*N_AB_*) to the number of times either A or B was observed (*N-D*, where *N* is the total number of sightings and *D* the number of times neither A nor B was seen) (66). Unless otherwise mentioned, data manipulation and analyses were carried out using MATLAB 7 (67).

### Social structure using networks

We examined social structure using network and cluster analyses. Social networks were constructed based on AI between individuals and visualized using Gephi 0.8.2 (68). The following network statistics were calculated: *degree* (the number of connections or *edges* arising from an individual or *node*), *clustering coefficient* (the proportion of all possible edges between the immediate neighbours of a focal node that actually exist, and, therefore, the probability that two randomly chosen neighbours of a focal individual are connected), *path length* (the number of edges on the shortest path between two nodes), and *distance-weighted reach* (the sum of the reciprocal of path lengths from a focal node to other nodes), calculated for individual nodes, and *density* (the proportion of all possible connections in the network that actually exist) and *modularity* (see below) calculated for the entire network (69-71). In order to find out whether the observed network was different from a random network, we compared the degree distribution of the observed network against Poisson expectation that would arise from an Erdös-Rényi random network (72). We also tested for preferred associations by randomly permuting the association data following Whitehead (73; Table S3). Network statistics of the Kabini 500-m dataset were compared with available statistics from previously studied populations. Since the mean and SD of these statistics were generally available from other populations, but the distributions were likely to be skewed and/or have different variances, we compared statistics across populations using Welch’s two-sample tests (74). It has been shown through simulations that the Welch’s test performs well under several scenarios involving the comparison of skewed distributions with unequal variances and sample sizes (75). As a further precaution, we used this test to compare statistics between the Uda Walawe and Samburu populations that had earlier been analyzed using randomization tests (24), and found the same results in all eight tests performed. Statistics from the Kabini 500-m dataset were used for comparison with the Uda Walawe and Samburu populations (as shown in 24).

Community structure within networks, and hence *modularity* (a measure of the extent to which a community is partitioned; this can be measured by comparing the fraction of edges within communities to that between communities), was identified using the Louvain method (43). The Louvain method identifies communities hierarchically and is known to be accurate. It uses a weighted network (in which edge weights, which are AI values between females, are incorporated rather than mere presence or absence of associations between females) in which each node is initially considered a separate community. Changes in modularity upon rearrangements of nodes are evaluated and rearrangements are stopped when a local maxima of modularity is obtained. The communities detected at this point are used as nodes for the next pass. Since the algorithm begins with rearrangements of single nodes across communities, this method does not suffer from the problem of identifying communities at a small scale. The algorithm is repeated iteratively until the maximum modularity is obtained, resulting in hierarchical partitions of communities within communities (43). This method allows for structure to be meaningfully examined at different hierarchical levels because the intermediate partitions correspond to local modularity maxima (43). This method, therefore, naturally lends itself to the investigation of social organization, when one is interested in finding out whether there are hierarchies or not. The Louvain method was implemented by calling the C++ codes made available by the authors (https://sites.google.com/site/findcommunities/) from MATLAB. We carried out the Louvain hierarchical community detection for the Uda Walawe and Samburu data also (data from 24, kindly provided by the authors) for comparison with the Kabini (Kabini 500-m dataset) population. We also constructed network structure curves following de Silva *et al.* (25) for comparison across populations. Here, the number of clusters (based on the Louvain method) with more than one female was plotted against AI threshold. The network structure curve provides information on the cohesiveness of the social network at different association strengths. Significant changes in the slope of the network structure curve were detected by comparing the values of number of clusters to the left and right of each point within a moving window of 0.3 using the Wilcoxon rank sum test (see 25).

### Effect of group size on AI and network statistics

We created random datasets, each with 100 individuals in 1500 sightings, distributed in group sizes following beta distributions with parameters that resulted in group size distributions that mimicked known elephant group size distributions (*α*=1, *β*=7, maximum group size=19 for Uda Walawe, *α*=2, *β*=9, maximum group size=26 for Samburu, and *a=1,* β=9.5, maximum group size=27 for the Kabini 500-m dataset; Kabini group size distribution from this study, Uda Walawe and Samburu group size distributions from 24). The maximum group size was altered to change average group size. We calculated the average, SD, and kurtosis of AI, and network statistics including average degree, average clustering coefficient, and average path length for the random datasets. One hundred random datasets were created for each beta distribution type with each maximum group size. Therefore, average group size and the AI or network statistics were averaged across these 100 replicates. We then plotted the statistic under consideration against average group size based on the random dataset, for each of the three beta distributions of group sizes, to visualize how the statistic changed with increasing average group size. For each observed statistic (calculated from field populations with an observed average group size), we calculated an expected random value of the statistic by interpolating the appropriate random curve (with matching beta distribution). Interpolation was done using cubic spline in CurveExpert version 1.37 (76). Using the 95% CI of the observed estimates, we calculated an interval with (expected - lower 95% CI of observed)/expected and (expected - higher 95% CI of observed)/expected values for each population. If these intervals overlapped across populations, it indicated that the populations differed from the random lines to the same extent and, therefore, differences in the statistic between the populations could be explained by differences in observed average group size. If the intervals of (E-O)/E did not overlap, it indicated that differences in the statistic between populations were significant beyond the effect of average group size. This was a conservative test because it was possible that the intervals of (E-O)/E could actually be larger than what we calculated based on 95% CIs.

### Cluster analysis and cumulative bifurcation curve using AI

Although hierarchical cluster analysis based on AI may not be useful for detecting hierarchical structure if social units at each tier of social structure show variability in AIs (also see 24), we used this method simply to compare the shapes of cumulative bifurcation curves across populations. We constructed dendrograms based on associations between individuals and used the plot of the cumulative number of bifurcations in the dendrogram at different linkage distances to identify knots (see 19). The average-linkage (UPGMA) method was chosen for clustering because it yielded the maximum cophenetic correlation coefficient value (CCC=0.976). Knots were identified by comparing the number of bifurcations in 0.2 and 0.3 windows above and below each point in the cumulative bifurcations plot, using a Wilcoxon rank sum test.

## Acknowledgements

This work was funded by the Department of Science and Technology’s (Government of India) Ramanujan Fellowship (to TNCV) under Grant No. SR/S2/RJN-25/2007 (dated 09/06/2008); Council of Scientific and Industrial Research, Government of India, under Grant No. 37(1375)/09/EMR-II and No. 37(1613)/13/EMR-II; National Geographic Society, USA, under Grant #8719-09 and #9378-13; and JNCASR. JNCASR provided logistic support. NS was given a Ph.D. scholarship by JNCASR, and PK was supported as a Ph.D. student by the Council of Scientific and Industrial Research. No. 09/733(0152)/2011-EMR-I. This work is part of the NS’s doctoral thesis. The funders had no role in study design, data collection and analysis, decision to publish, or preparation of the manuscript.

We thank the office of the PCCF, Karnataka Forest Department, for field permits. We are grateful to various officials of the forest department, from the PCCF and APCCF, to the Conservators of Forests and Range Forest Officers, and the staff of Nagarahole and Bandipur National Parks for permits and support at the field site. Deepika Prasad, Arjun Ghosh, and Hansraj Gautam helped with field data collection. Mr. Krishna, Mr. Gunda, Mr. Althaf and others provided field assistance. We thank TNC Anand, Indian Institute of Technology Madras, for help with the community detection codes, and Ajay Desai, WWF, for useful discussions.

